# A guided approach for subtomogram averaging of challenging macromolecular assemblies

**DOI:** 10.1101/2020.02.01.930297

**Authors:** Danielle Grotjahn, Saikat Chowdhury, Gabriel C. Lander

## Abstract

Cryo-electron tomography is a powerful biophysical technique enabling three-dimensional visualization of complex biological systems. Macromolecular targets of interest identified within cryo-tomograms can be computationally extracted, aligned, and averaged to produce a better-resolved structure through a process called subtomogram averaging (STA). However, accurate alignment of macromolecular machines that exhibit extreme structural heterogeneity and conformational flexibility remains a significant challenge with conventional STA approaches. To expand the applicability of STA to a broader range of pleomorphic complexes, we developed a user-guided, focused refinement approach that can be incorporated into the standard STA workflow to facilitate the robust alignment of particularly challenging samples. We demonstrate that it is possible to align visually recognizable portions of multi-subunit complexes by providing *a priori* information regarding their relative orientations within cryo-tomograms, and describe how this strategy was applied to successfully elucidate the first three-dimensional structure of the dynein-dynactin motor protein complex bound to microtubules. Our approach expands the application of STA for solving a more diverse range of heterogeneous biological structures, and establishes a conceptual framework for the development of automated strategies to deconvolve the complexity of crowded cellular environments and improve in situ structure determination technologies.

## Introduction

Cryo-electron microscopy (cryo-EM) is an impactful methodology for three-dimensional (3D) structural determination of macromolecular complexes. While single particle EM gained widespread notoriety for its utility in solving high resolution structures of purified proteins, cryo-electron tomography (cryo-ET) has emerged as the leading technique for visualizing the structures of large, transient, dynamic, flexible, and/or heterogeneous samples in native or near-native reconstituted cellular environments^1,2^. The implementation of automated data collection^3^ packages^4,5^ and optimized tomographic acquisition schemes^6–9^, combined with direct electron detectors, energy filters, and phase plates^10^ has revolutionized the feasibility of visualizing cellular machinery for functional and physiological interpretation^11^. Multiple copies of the biological complex of interest can be identified within reconstructed tomograms, and 3D “subvolumes” can be extracted and averaged together in a process called subtomogram averaging (STA)^12–20^ to obtain better resolved 3D reconstructions of the complex of interest. Notably, aspects of single-particle image processing have been incorporated into STA processing packages^18,19,21^, and combined with improved 3D-contrast transfer function (CTF) estimation and missing-wedge compensation^20,22–24^to achieve reconstructions in the sub-nanometer resolution regime^9,20,25–39^. Notably, high-resolution (3.9-3.1 Å) reconstructions of the HIV-1 capsid-SPI region of the viral Gag protein has been determined using cryo-ET^20,23,40^, further emphasizing the promise of this technique in obtaining high-resolution structural information of complexes *in situ*.

However, despite improvements in instrumentation and algorithms, the field is still far from routinely obtaining high resolution structures by STA, as most structures deposited in the EM Data Bank (EMDB), and determined by this method are at resolutions worse than ~20 Å (**Figure 1a**). Moreover, while the ability to elucidate the structures of pleomorphic, multi-subunit complexes in native, *in situ* cellular environment is a major advantage of cryo-ET and STA over other structural techniques, current STA processing strategies are typically only successful for highly ordered, symmetrical, homogenous samples that have limited conformational variations, and are present in high copy numbers within a single tomogram (**Figure 1b**). Examples of such complexes include purified viruses and associated viral complexes^17,25,27–29,31,33,37–52^, highly-abundant cytoplasmic and membrane-associated ribosomes^9,10,19,35,53–62^, and axonemal dynein motors^63–83^. Large, eukaryotic and bacterial membrane-associated complexes have also benefited greatly from this technique, as alignment of the relatively high-signal membrane bilayer can help drive the initial alignment of the more noisy, low SNR complex of interest. Examples of these complexes include COPI/II vesicle coats^34,84,85^, bacterial secretion systems and flagellar motors^86–98^.

**Figure 1.**
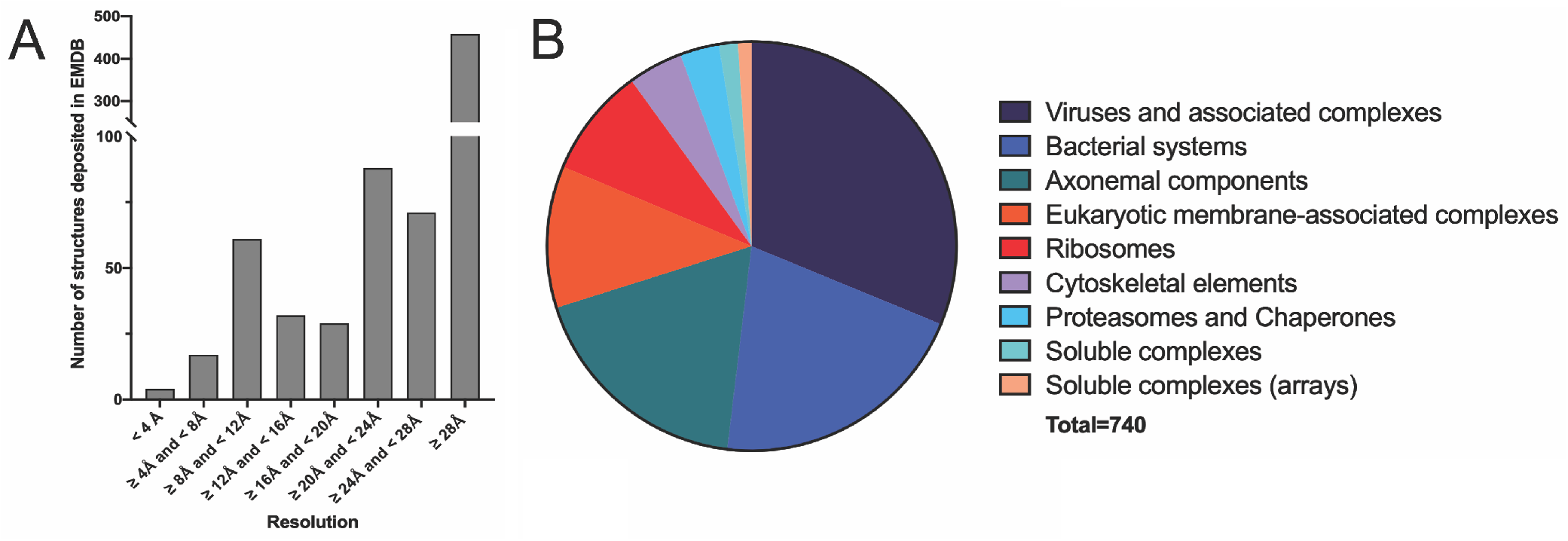
Subtomogram averaging: current state of the field. **A.** Histogram displaying resolution distribution of structures solved by subtomogram averaging. While some high resolution (<4 Å) structures have been reported, the majority of the deposited structures solved by subtomogram averaging are low resolution maps. **B.** Pie chart demonstrated the types of biological complexes solved by subtomogram averaging. For both (A) and (B), data was downloaded from Electron Microscopy Data Base (EMDB) in October 2019 and sorted by resolution and biological complex, respectively.

A major challenge facing STA of macromolecular machines embedded in crowded environments is in dealing with the high degree of compositional heterogeneity present within each individual subvolume. Ideal extracted subvolumes should only contain signal from the complex of interest. However, in most cases, the extracted subvolumes contain a variety of biological macromolecules in addition to the specific macromolecule of interest. This is particularly relevant for tomograms of cellular landscapes teeming densely packed macromolecules, which may include but not limited to signal-dominant membrane structures, large and featureful filaments, or an abundance of cytosolic complexes. The presence of a diverse array of different subcellular structures, each with their own signature electron scattering profile, often leads to misalignment of the targeted complex when subjected to even the most sophisticated STA algorithms for 3D classification or refinement. Therefore, one of the fundamental areas for growth for these methodologies is the development of processing strategies to deal with high levels of heterogeneity present within individual, low-signal subvolumes, and successfully generate 3D structures of dynamic, heterogeneous complexes that may only be present in relatively small numbers within tightly-packed cellular or near-native reconstituted systems.

In order to address some of these challenges inherent to heterogeneous samples, we developed a user-guided, focused refinement approach to elucidate 3D structures of large, flexible, non-symmetric biological complexes present in relatively low abundance within individual tomograms. This user-guided approach has the potential to overcome several limitations described above, and will be applicable to solve structures of macromolecular complexes that display recognizable, known features that are discernible to the user within the subvolumes, but not identified by automated particle selection programs, and/or not reliably aligned by 3D classification or refinement algorithms. Users may encounter such situations when the biological target is (1) present within a “crowded” subcellular environment, where many diverse and variable biological features obfuscate the target; (2) dynamic and/or flexible in such a way that every targeted complex represents a unique conformational species; and/or (3) associated with another large, high-signal biological complex, and initial attempts using STA result in alignment of this signal-dominant feature instead of the targeted complex.

We describe here the overall methodology and rationale for the user-guided, focused refinement approach. We also demonstrate how this strategy overcame significant challenges and was successfully applied for elucidating the first 3D reconstruction of the large, flexible, asymmetric microtubule (MT)-bound dynein-dynactin-BicaudalD2 (DDB-MT) complex^99^. Although this methodology involves manual user input, we demonstrate that it is free of reference model or user bias.

## Materials and Methods

### Subvolume Extraction

The first step involves manually identifying the complexes within the reconstructed tomogram for 3D subvolume extraction. This process can be performed using any of several STA processing packages, such as EMAN2^18^, PEET^14,100^, Dynamo^16,101^ or PyTom^15^ (**Figure 2a**). For ease in identification of complexes of interest in subsequent steps in this procedure, it is recommended that particles are initially extracted from tomograms reconstructed using iterative reconstruction methods such as Simultaneous Iterative Reconstruction technique (SIRT)^102^ or Simultaneous Algebraic Reconstruction Technique (SART)^103^. A list of coordinates denoting the approximate centroid of the identified complexes should be generated and used to extract subvolumes from each tomogram, so that the complex of interest is roughly positioned in the center of the extracted volume. The box size of the extracted subvolumes should be sufficient to accommodates the entirety of the complex of interest, but limit the inclusion of significant off-target biological material.

**Figure 2.**
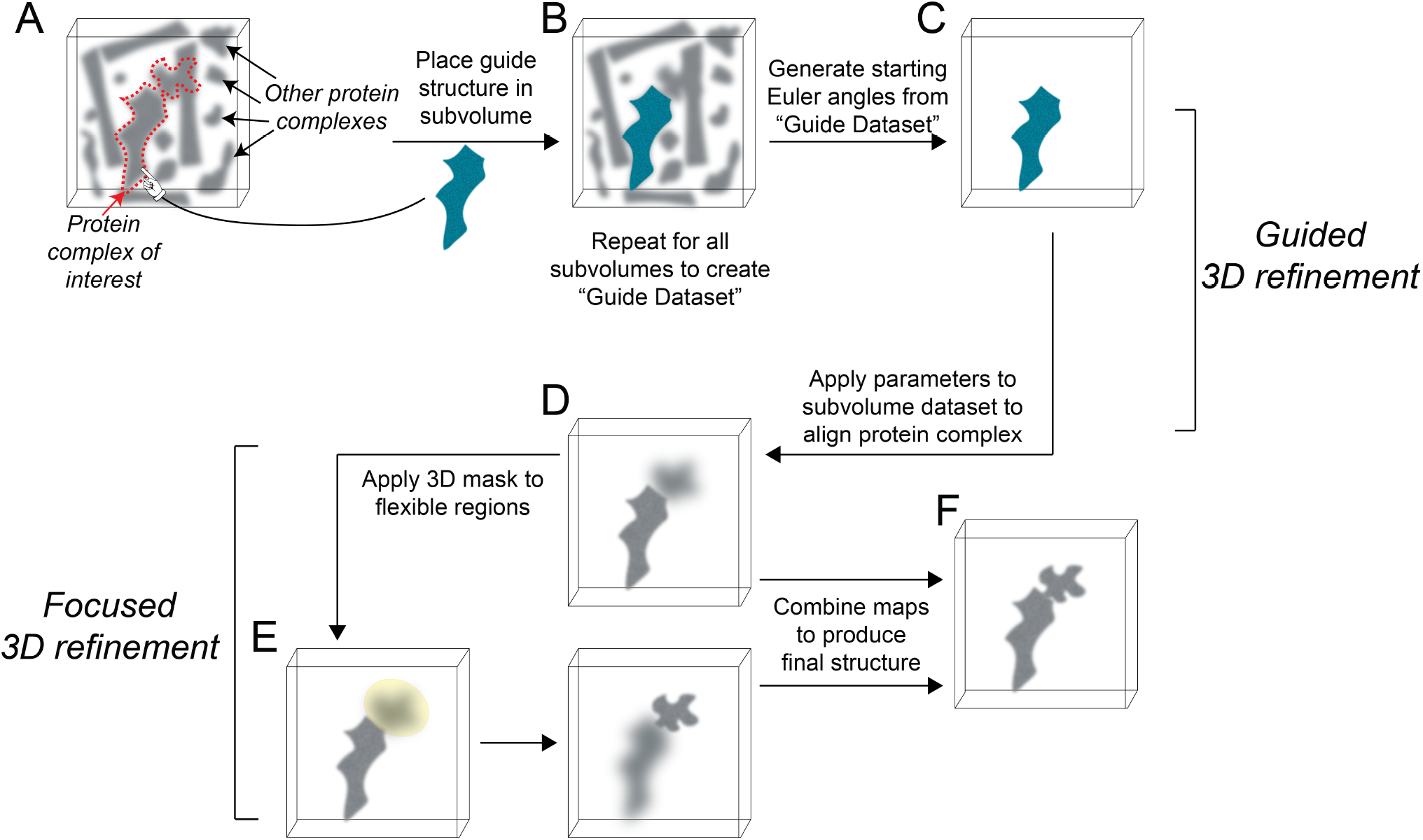
Workflow for guided subtomogram averaging approach. **A.** Cartoon representation of an extracted subvolume from a noisy tomogram. The biological complex of interest is outlined by red dashed line. **B.** A reference structure, shown in cyan, of a portion of the complex is manually docked into the subvolume in the same location as it is presented in the original, extracted subvolume. **C.** These docked reference structures are saved as individual subvolumes, creating a “guide dataset” of reference structures that represent the positions of these complexes in the original subvolume. **D.** An initial alignment of the “guide dataset” is performed, and the Euler angles assigned to the reference structure during alignment are applied to particles from the original, extracted subvolume dataset to yield a final, aligned structure of the complex of interest. **E.** To resolve the highly flexible regions, a soft, 3D binary mask (shown in yellow) can be used to focus the alignment to these regions. **F.** Individual focused maps can be combined to produce the final, well aligned structure.

### Identification of targeted complexes within individual subvolumes

Once the 3D subvolumes have been extracted, the user must assign an approximate 3D orientation (x, y, z, phi, theta, psi) to the complex located at the center of the subvolume. This can be performed by simultaneously viewing (1) an extracted subvolume, and (2) fitting a 3D map of a known portion of the complex (referred to herein as “guide structure”), either simulated from an independently determined crystal structure or EM reconstruction, within a visualization program, such as UCSF chimera^104^ (**Figure 2a**). It is recommended that the user initially manually orients the guide structure to match the location of the corresponding feature of interest in the extracted, noisy subvolume, and then uses a cross-correlation-based fitting program (such as UCSF Chimera’s “fit in map”) to more precisely dock the reference structure into the local feature of interest within the subvolume (**Figure 2b**). Often, alternating the viewing mode from 2D planes/slices of the subvolume to 3D isosurface rendering within UCSF Chimera can aid in identifying the features of interest and placing the guide structure within the extracted subvolume. The “docked” 3D guide structure should be assigned the coordinate system, voxel size and bounding box dimensions of the originally extracted subvolume by using the “vop resample” command in UCSF Chimera, prior to saving it as a new volume. To facilitate downstream processing, a naming system that contains the original extracted subvolume should be used when saving the docked structure. Alternatively, if the translational and rotational parameters associated with the docked 3D structure can be saved in a format that is compatible with an image processing package such as RELION, a refinement of the raw, extracted subvolumes can now immediately be performed using a local search based on the assigned orientations (skip the next step).

### Initial refinement using docked structures

Each of the originally extracted subvolumes now have a corresponding “guide” subvolume from the docked guide structure, which denotes the 3D orientation of the targeted biological complex within the original subvolume. This “guide dataset” is now used as input for a 3D refinement/reconstruction program such as RELION auto-refine in order to retrieve the rotational and translational parameters of the docked features of interest (**Figure 2c**). These parameters will serve as the initial alignment parameters for local refinement of the feature of interest in the originally extracted volumes, using a subtomogram averaging program (i.e. RELION). Given the high SNR of the guide dataset (essentially SNR=1.0), the 3D refinement of the guide dataset should converge to a reconstruction that is identical to the 3D reference structure used for docking (**Figure 2c**).

### Local refinement using raw, extracted subvolumes

The rotational and translational parameters derived from alignment of the guide dataset to a common reference are now applied to the raw, extracted subvolumes. The user can edit the metadata file containing these parameters for the guide dataset subvolume, replacing the path and name of each guide volume with that of the corresponding originally extracted subvolume. For example, if using RELION, one would replace the filenames in the “rlnXXX” column of the data.star file with the names of the originally extracted volumes. These orientation parameters serve as a starting point for a local orientational search. Given the low SNR of the experimental data, it is important that this search and translation range not be too generous, as the refinement may diverge substantially from the targeted complex. For example, In RELION, this could be accomplished by performing a 3D auto-refinement as a continuation of the previous alignment, or by setting the initial angular sampling and local searches from auto-sampling to the same small increment. A successful 3D refinement will converge to a reconstruction that contains recognizable structural features of the complex of interest that are generally similar, but not identical to those that were marked previously by the 3D reference volume (**Figure 2d**).

### 3D focused refinement of flexible regions belonging to the targeted complex

If the alignment procedure described above leads to a well-resolved structure of the entire biological complex of interest, then no further processing is needed. However, in most cases, this STA processing strategy will be applied to macromolecular complexes that are larger than the guide structure used to assign orientations. Notably, the complexes targeted using this approach are likely challenging for automated methods due to the presence of flexible regions, which will likely not be resolved in the reconstructed density. However, since a portion of the complex in each subtomogram has now been assigned a common orientation, an attempt can be made for refining these flexible portions, using a 3D binary, soft-edged masks (ellipsoidal or spherical) corresponding to individual sub-regions of the complex (**Figure 2e**).

### Combining results of individual focused refinements to produce a single 3D structure of the macromolecular complex of interest

The individual maps resulting from focused 3D alignment and averaging of each sub-region of the biological complex can be combined using the “vop maximum” function in UCSF chimera which measures and retains the maximum voxel values of overlapping volumes to produce a final composite reconstruction (**Figure 2f**).

## Results and Discussion

### Challenges to working with the microtubule-bound dynein-dynactin-BicaudalD2 (DDB) complex: a sample dominated by variability, heterogeneity and flexibility

Due to its inherent asymmetry, relatively large size (~1.5 MDa) and multi-subunit complexity (**Figure 3a**), cytoplasmic dynein represents a challenging target for 3D structure determination. Dynein is activated for microtubule-based transport upon binding to additional cofactors, including another large, (~1.5 MDa) multi-subunit complex called dynactin (**Figure 3a**). In order to investigate how these cofactors activate dynein for microtubule-based transport, we developed a near-native reconstitution system to purify microtubule-bound dynein-dynactin complexes from mouse brain lysate and performed cryo-ET to visualize the 3D architecture of this transport complex^99^. Although the complexes were readily visible within reconstructed 3D tomograms (**Figure 3b**), there were several unique challenges that prevented the application of automated, template-based approaches for subvolume selection. For example, due to slight variations in the amount of endogenous proteins (tubulin, dynein, dynactin), both the total number and the binding pattern of DDB complexes on individual microtubules varied dramatically between sample preparations and within tomograms derived from the same batch of prepared sample (**Figure 3b**). In order to ensure sufficient number of DDB complexes per tomogram, for downstream processing by STA, we increased the concentration of decorated microtubules deposited on the EM grid, thus increasing the number of DDB complexes per tomogram. However, this also led to an increase in the overall ice thickness of the sample, further decreasing the SNR of individual extracted subvolumes, wherein the signal from the microtubule densities dominate, and obfuscate the sparsely decorated dynein-dynactin complexes (**Figure 3c**). These challenges precluded our ability to use of automated search algorithms to select DDB particles for 3D subvolume extraction, including those programs that utilize filament models to define the particle orientation relative to a tubular structure^101^.

**Figure 3.**
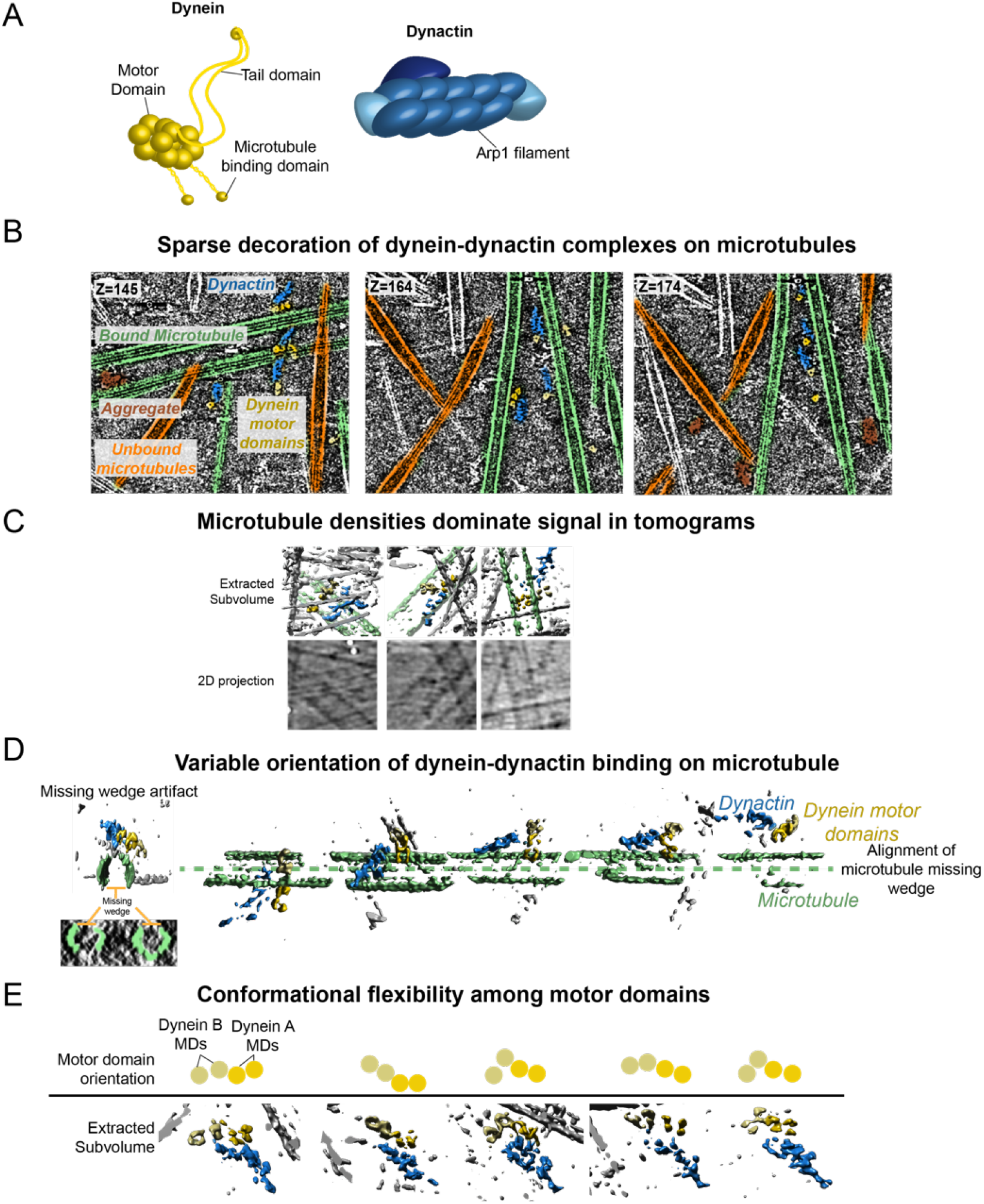
Microtubule-bound dynein-dynactin complex: a challenging structural target. **A.** Cartoon representations of cytoplasmic dynein (left, yellow), and dynactin (right, blue) with labeled components of major structural and functional domains. **B.** Representative X-Y slices of a reconstructed, three-dimensional tomogram progressing through the z-axis (z=145, 164, 174; from left to right). Each tomographic slice is colored to indicate different components present within in vitro reconstituted dynein-dynactin transport environment, included dynein motor domains (yellow) in complex with dynactin (blue), bound to microtubules (green), as well as non-specific protein aggregates (brown) and microtubules that appear devoid of bound dynein-dynactin complexes (orange). **C.** Three representative extracted subtomograms and corresponding 2D projections illustrate how the microtubule densities present in each individual subvolume can dominate the signal, likely making it difficult for computational algorithms to automatically extract and align voxels containing the relatively lower-signal density of the dynein-dynactin complexes. **D.** (left) Representative extracted subtomogram with components colored same as (B), displaying the missing wedge density artifacts observed in tubular microtubule structures. **D.** (right) Five representative extracted subtomograms displayed with aligned microtubule missing wedge, showing the degree of variability that the dynein-dynactin complex can bind to the microtubules when taking into account alignment of the missing wedge of the microtubule. **E.** Five representative extracted subtomograms (bottom) colored similar to (B), oriented such that the dynactin density (blue) is in the same position for each. Colored circles (top) represent position of motor domains in the corresponding subtomogram, and are used to illustrate the range and diversity of conformational flexibility among the four dynein motor domains that is unique to each individual subtomogram.

We attempted multiple strategies to align, classify and average the subvolumes using RELION subtomogram averaging workflow (**Figure 4**). However, none of our initial attempts were successful at producing a well-resolved structure with features that resembled previously-resolved portions of the microtubule-bound dynein-dynactin complex^105,106^. In most instances, these programs produced a structure that resembled a microtubule in close proximity to an uninterpretable density that might correspond to DDB complexes, however there was no clear indication of dynein’s characteristic structural features, including the donut-shaped motor domains, or dynactin’s arp1 filament (compare cartoon diagrams in **Figure 3a** with structures in **Figure 4a**). Strategies to eliminate the putative microtubule density using a 3D soft mask to focus refinement to this putative DDB complex were similarly unsuccessful (**Figure 4b**).

**Figure 4.**
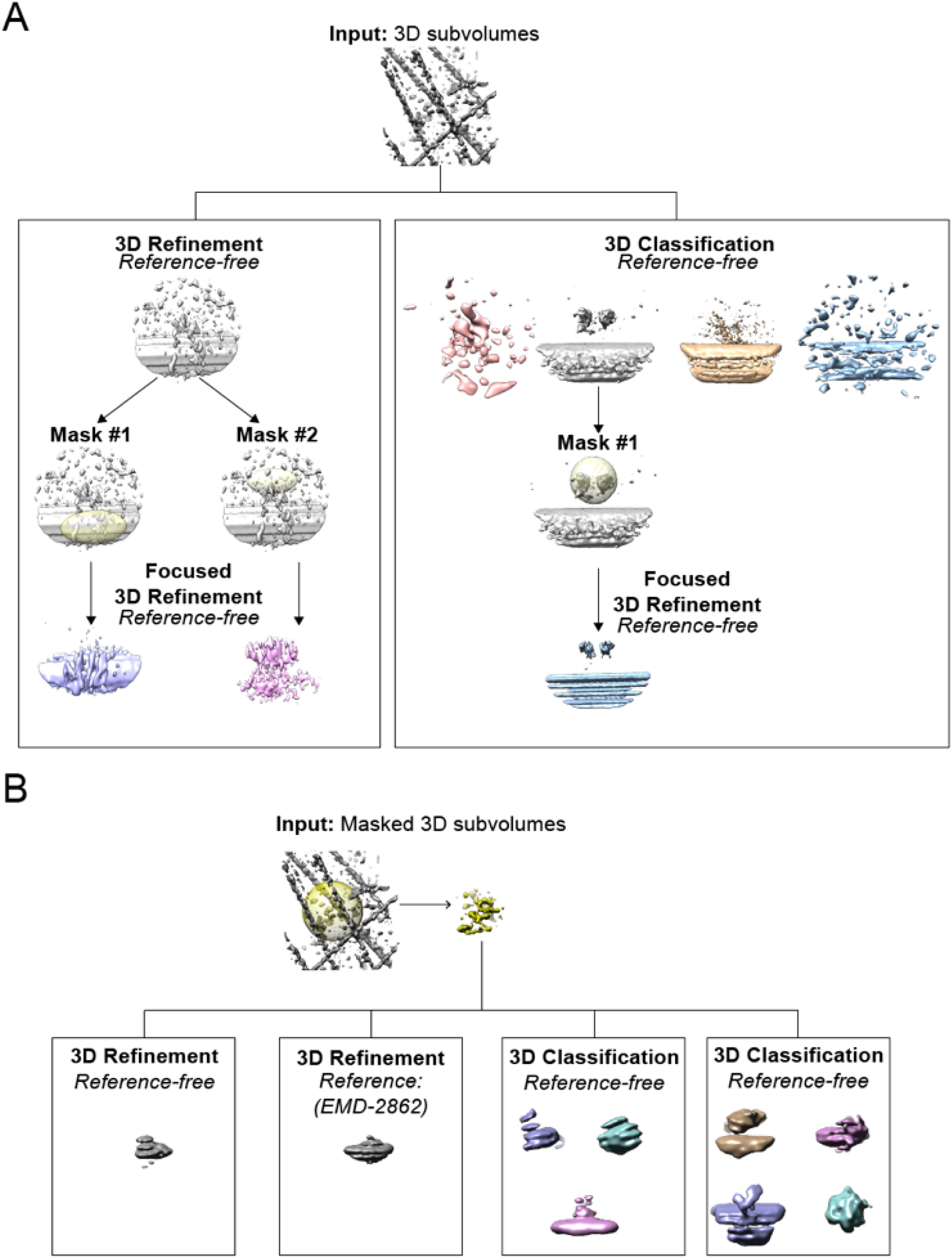
Unsuccessful attempts at structure elucidation of dynein-dynactin complexes using conventional subtomogram averaging approaches in RELION. **A.** Data processing strategy using 500 extracted subvolumes containing dynein-dynactin complexes as input. Reference-free 3D refinement, 3D classification and focused 3D refinement strategies failed to produce a well-aligned structure with characteristic features that resembled the dynein-dynactin complex. **B.** Data processing strategy using masked 3D subvolumes as input to reduce misalignment of other densities that are present in larger subvolumes in (A). Simliar to (A), both 3D refinement and 3D classification failed to produce a well-aligned structure with the masked subvolumes as input.

These inconclusive results were surprising, since manual inspection of X-Y slices of tomograms demonstrated clearly discernible structural features corresponding to dynactin and dynein complexes (Figure 3b). We posit that that several unique features of the DDB-MT complex sample can explain this discrepancy. For one, the “missing wedge” of information that results from the inability to fully capture all tilted views of the sample during tomographic image acquisition is most notable in the microtubule filaments within extracted subvolumes, where missing 2D projections result in large sections of microtubules lacking density (**Figure 4d**). Averaging of extracted subvolumes is likely driven by the alignment of the missing wedge artifact within microtubule densities, rather than by the comparably smaller, lower-signal DDB complexes (**Figure 4 c & d
**). Alignment of the microtubule missing wedge results in significant misalignment of DDB complex, which is likely exacerbated by the variability by which the DDB complexes orient on the microtubule (**Figure 4d**), and further compounded by the extreme flexibility among the motor domains within individual DDB complexes (**Figure 4e**).

As mentioned previously, to overcome the sparse decoration of dynein-dynactin complexes on microtubules, we increased the overall concentration of sample deposited on the EM grid, leading to a crowding of microtubule structures within tomograms. In addition to impeding our ability to utilize automated template matching for particle picking (see above), the crowded nature of the sample often resulted in extracted subvolumes that included a single dynein-dynactin complexes surrounded by numerous microtubules, which likely also contributed to mis-alignment observed in our initial STA attempts. Although we attempted to extract with a box size that minimally included the full, dynein-dynactin complex while excluding neighboring microtubule densities, the extracted subvolumes nonetheless often included many structures in addition to the desired dynein-dynactin complex of interest (**Figures 3b &d**). In summary, despite many diverse attempts and strategies, the unique features of the DDB complex regarding variability in sample preparation, missing wedge alignment of signal-dominant microtubules, and extreme heterogeneity within the DDB complex itself all likely contributed to the difficulty in obtaining an interpretable 3D DDB-MT structure using traditional STA methodologies.

### Manual docking of reference dynein tail-dynactin-BicaudalD2 structure into raw subvolumes

In order to provide *a priori* information to guide the alignment of microtubule-bound dynein dynactin complexes, we visualized individual 3D subvolumes extracted from tomograms reconstructed by SIRT in UCSF Chimera. To reduce noise and facilitate the identification of the characteristic features of the DDB complex (**Figure 5a & b**), a Gaussian filter with a width of 8.52 Å was applied to the subvolumes using the “volume filter” function in UCSF chimera. Once identified, a structure of the dynein tail-dynactin-BicaudalD2 complex map (EMD-2860, hereafter referred to as TDB) was opened in the same chimera session, resampled on the same grid as the extracted 3D subvolumes, repositioned to match similar features within the extracted subvolume, and the Chimera “fit in map” function was used for more precise fitting (**Figure 5b**). Once the EMD-2860 structures were successfully positioned, the docked EMD-2860 guide structure was saved in position within the same 3D grid as the original subvolumes (**Figure 5c**). Thus, the 3D reference now reflected the observed position of the reconstructed complex within the subvolume. This manual positioning of the TDB reference structure was determined to be an optimal strategy since attempts to automate the fit-in-map function were largely unsuccessful, often leading to the incorrect positioning of the TDB structure within a high signal component within the subvolume, including the microtubule or protein aggregate (**Figure 6**). This process was repeated for all 502 individual, extracted subvolumes containing microtubule-bound dynein-dynactin complexes.

**Figure 5.**
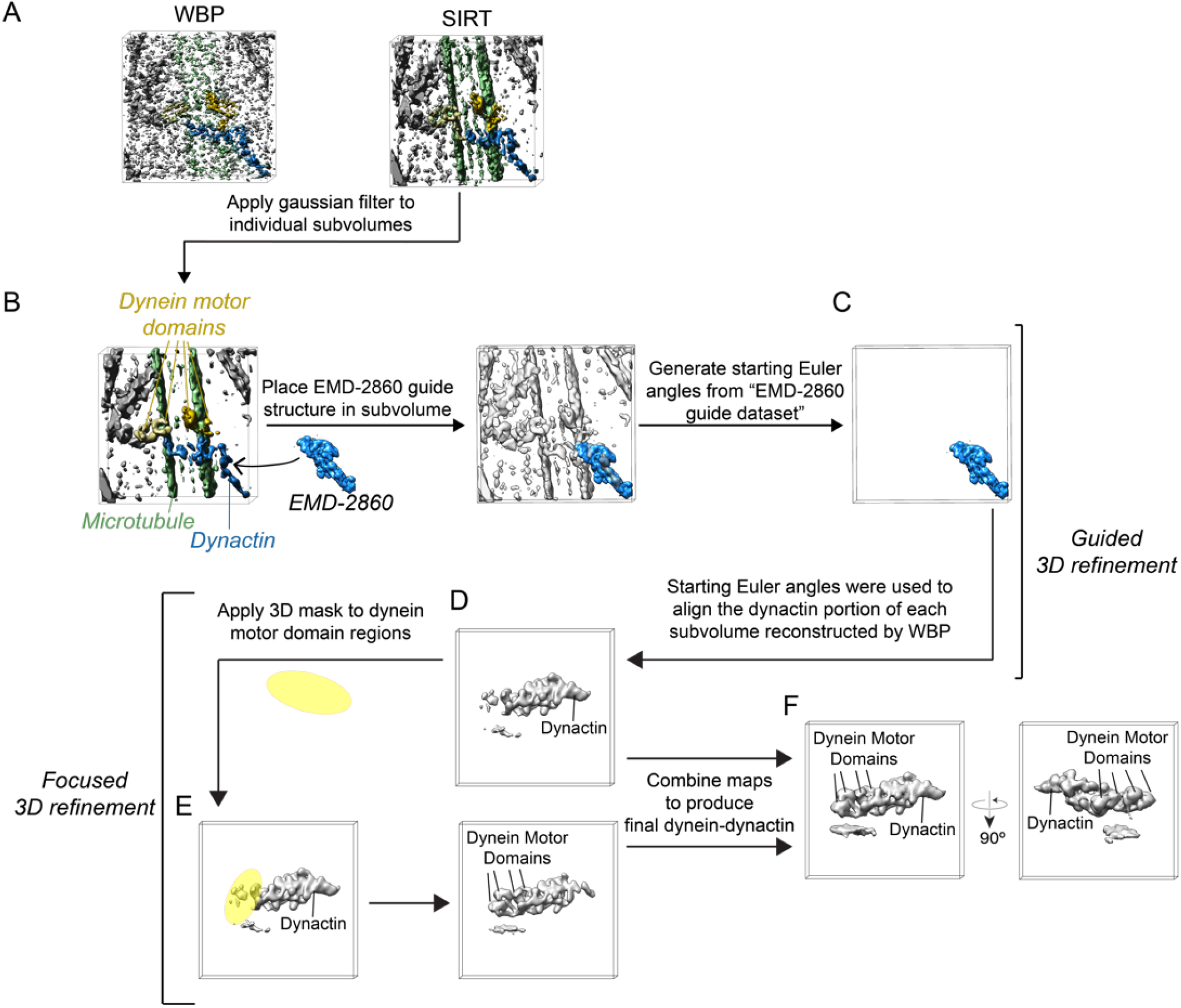
Application of guided approach to solve structure of dynein-dynactin complex. **A.** Comparison of subtomogram extracted from tomogram reconstructed by weighted back projection (WBP, left) versus simultaneous iterative refinement technique (SIRT, right). **B.** To further enhance clarity of dynein-dynactin features present within subvolumes, extracted subvolumes from SIRT reconstructions were Gaussian filtered. A cryo-EM map of the dynactin guide structure (EMD-2860) was docked into SIRT reconstructed, Gaussian filtered subtomograms using UCSF Chimera. This process was repeated for ~500 subvolumes to generate the “EMD-2860 guide dataset.” In (A) and (B), portions of the subvolumes are colored to indicate location of microtubule-bound dynein-dynactin complexes (dynein motor domains, yellow; dynactin, blue; microtubule, green). **C.** Alignment of the “EMD-2860 guide dataset” was performed using RELION 3D autorefine to generate starting Euler angles. **D.** The starting Euler angles generated in (C) were applied to subvolumes extracted from tomograms reconstructed by WBP to align the dynactin portion of the dynein-dynactin complex. **E.** 3D ellipsoidal, soft-edge binary masks (yellow) were used to perform a focused 3D refinement to better resolve the flexible dynein motor domains. **F.** The two maps generated from the initial alignment of dynactin (D) and the focused 3D refinement of the motor domains (E) were aligned and stitched together to generate a combined structure of the dynein-dynactin complex.

**Figure 6.**
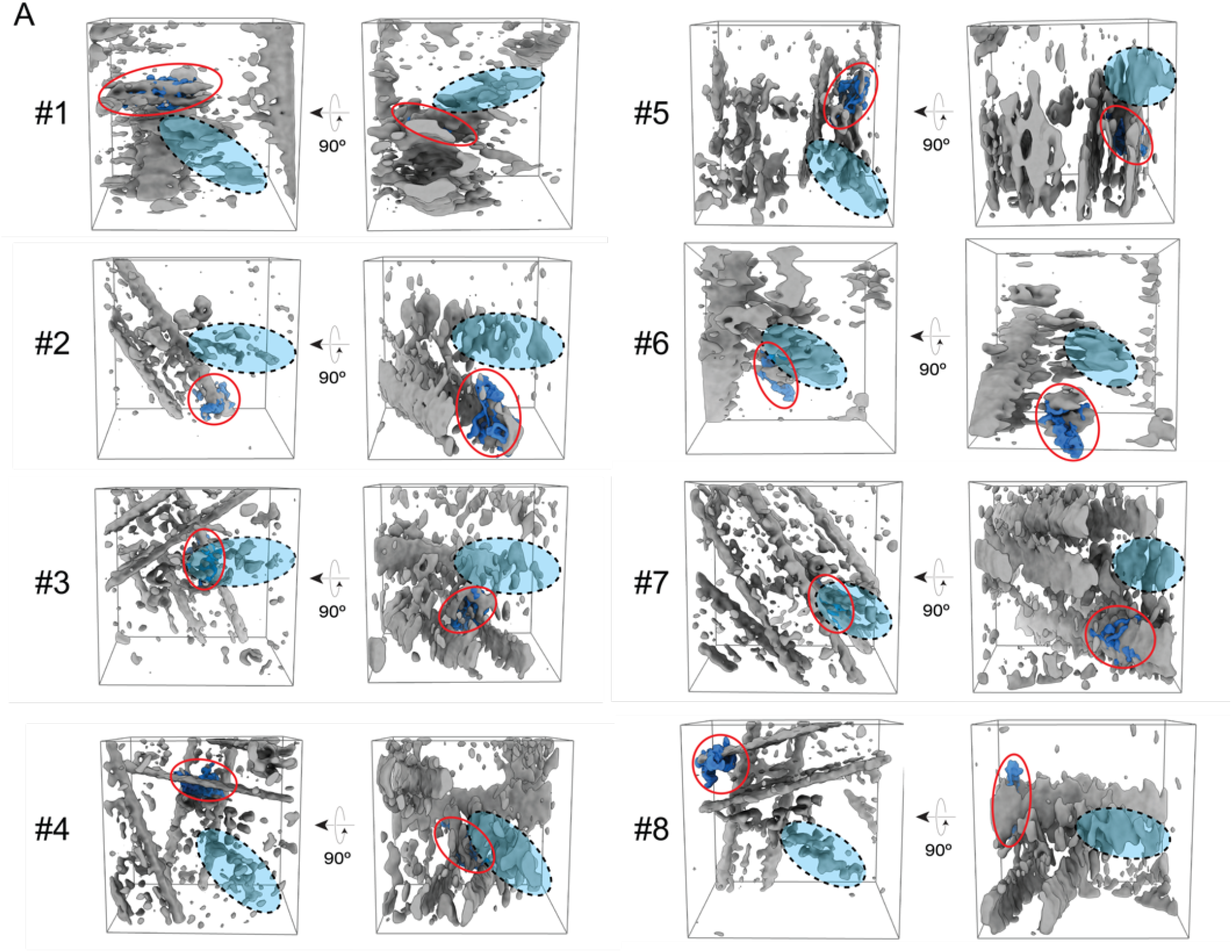
Failed attempts to automate docking of reference structure. **A.** Representative subvolumes (#1-8, in gray) with the position of the dynactin portion of the dynein-dynactin complex highlighted with a light blue oval outlined in black dashed lines. Red circles indicate the position of the guide structure (EMD-2860, dark blue) with the highest cross-correlation score docked in the subvolume, as determined by the UCSF Chimera “Fit in map” program. In all cases, the position with the highest cross-correlation score (red circle) does not overlap with the correct position of dynactin within the guide structure (light blue oval outlined in black dashed lines), demonstrating the difficulty in utilizing automated approaches to manually dock in guide structures into extracted subvolumes of the dynein-dynactin complexes. Due to this low success rate of correctly docked structures using this approach, initial manual docking of reference structure into extracted subvolumes was used for successful execution of the guided approach for dynein-dynactin complex.

### 3D alignment of guide (EMD-2860) structure

The guide dataset was used as input into RELION 3D autorefine, and after approximately 40 iterations the refinement converged, as expected, to a single structure identical to the low-pass filtered EMD-2860 guide structure. The calculated angular and translational parameters from the initial alignment were applied to the corresponding subvolumes from the original dataset, and a single iteration of 3D auto-refinement in RELION lead to a structure that resembled the TDB reference, with the notable addition of an extra dynein tail-like density projecting from the dynactin-BicaudalD2 scaffold. Furthermore, additional densities likely corresponding to the flexible dynein motor domains could be observed in the reconstruction (**Figure 5d**). Presence of these additional densities, that were missing in the guide structures, forms internal controls that confirm the final reconstructed volume to be free from any bias imposed by the guide structure. To further resolve these flexible regions, we used a 3D soft-edge binary mask that surrounded these portions of the complex and perform a refinement using local angular and translational searches (**Figure 5e**). The resulting well resolved sub-structures were fitted relative to each other and then combined into a composite 3D structure of the microtubule-bound DDB complex^99^, using the “vop maximum” function in UCSF chimera (**Figure 5f**).

### Control experiments to test for user and reference bias

As noted in previous studies, the presence of additional, physiologically-relevant densities within our aligned structure, including an additional tail density bound to dynactin that is absent in the original reference TDB structure, as well as the appearance of four motor domain densities, all serve as internal, positive controls that suggest there is no user or reference bias influencing the final 3D structure (**Figure 5f**)^99^. Furthermore, all 3D refinement jobs were performed without using an initial starting model for alignment, thus preventing the introduction of reference bias during the initial iterations of refinement. However, due to the use of user-generated input as a guide model for this approach, we performed a control experiment that can specifically test for bias introduced by user or reference, by randomly assigning starting Euler angles to the raw particles. In three separate experiments, this random assignment of starting Euler angles caused complete misalignment of the final 3D structure (**Figure 7**). This control experiments suggest that the success of this user-guided approach critically depends on the correct and accurate identification of the feature of interest within the crowded, low-signal subvolumes, and that the reference structure used as no influence on the reconstruction of the final, 3D structure.

**Figure 7.**
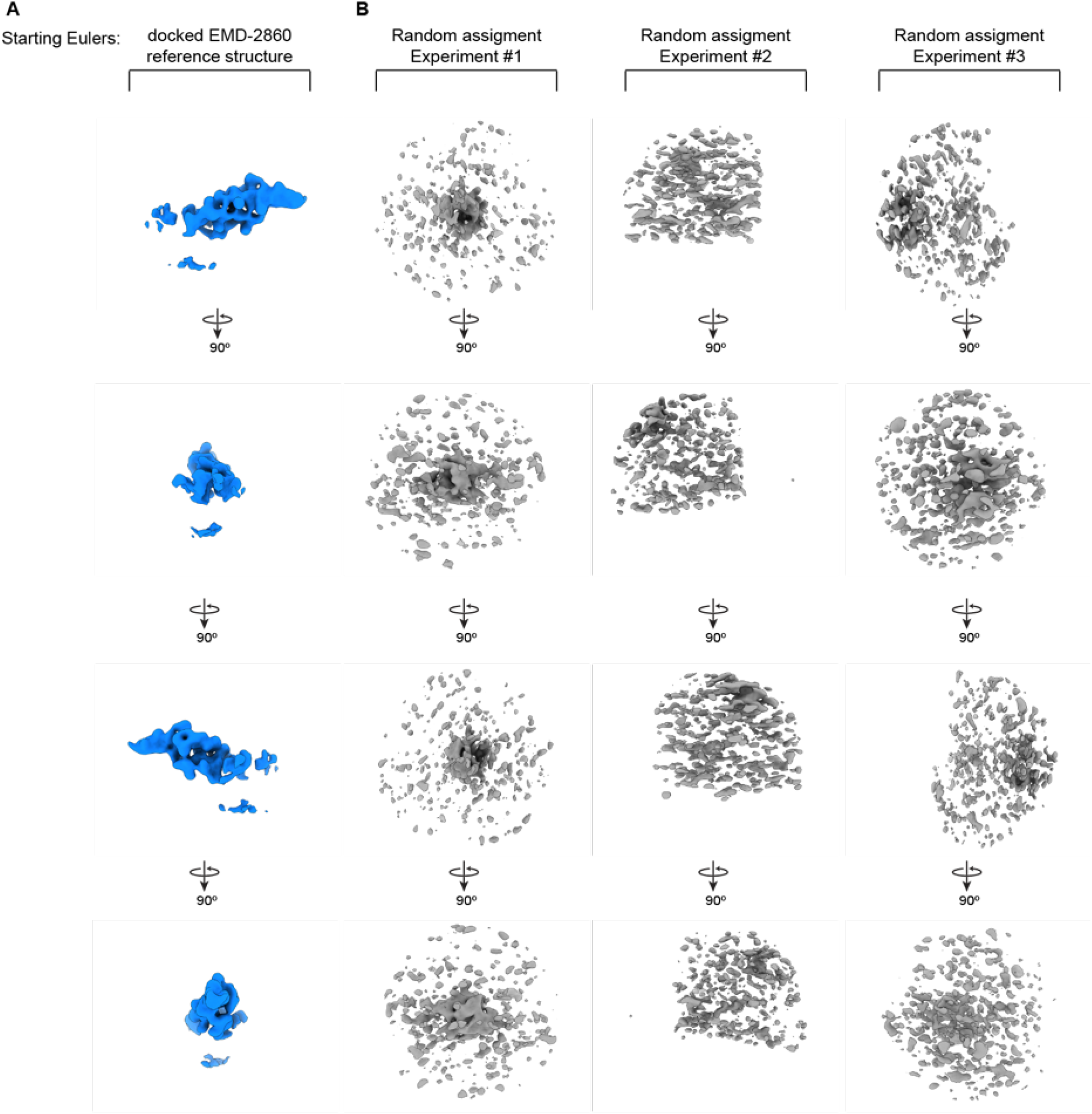
Control experiments to test for model bias in guided subtomogram averaging procedure. **A.** The resulting converged density that corresponds to the dynactin portion of the dynein-dynactin complex when the starting Eulers are assigned using the guided subtomogram averaging procedure using the manually-docked dynactin reference density (EMDB 2860) in the sub-volumes of dynein-dynactin complexes. **B.** The resulting densities from three distinct experiments generated from random assignment of the starting Eulers for the dynactin complex using three distinct starting seed models. The lack of any discernable features resembling the dynein-dynactin complex in these three control experiments suggest that the guided subtomogram averaging strategy does not introduce any model bias or other artifacts to the final subtomogram averages of the dynein-dynactin complex.

## Concluding Remarks

Despite significant advancement in 3D classification and refinement algorithms, the accurate alignment of subcellular structures using STA still remains a challenging endeavor. In many cases, the human eye can more readily identify certain 3D objects and features within noisy subvolumes than even the most advanced computational algorithms. For this reason, segmentation or annotation of features within tomograms is still typically done in a manual fashion within the cryo-ET field using programs such as IMOD^107^. The approach described here takes advantage of the visual expertise of the user to tease out the location of the biological complex of interest, and uses this information to help guide subtomogram averaging algorithms to align common features present within hundreds of individual subvolumes. We show that it is possible to preferentially align low-signal features (i.e. dynein-dynactin complex) relative to signal-dominant structures (i.e. MTs) by simply providing *a priori* information to help “guide” the refinement using RELION auto-refine. The success of this approach to align biological assembly within a complex, near-native reconstituted system suggests that this methodology has the potential to be successful for *in situ* structure determination of visually recognizable macromolecules present within crowded, cellular environments.

While the manual identification of the complex of interest within hundreds of individual subtomograms is user-intensive and laborious, for particularly challenging samples, this strategy may be the only feasible method for structure determination, especially in cases where prior attempts using traditional STA programs have failed. The basis of this approach in guiding alignment procedures to identify specific features within noisy subvolumes could eventually become an automated procedure using machine-learning algorithms^108^ or crowd-sourcing approaches^109^, thus reducing the amount of user-input. However, to our knowledge this strategy is the only method that has been shown to work for incredibly challenging macromolecular complexes using currently available computational processing tools and algorithms.

## Acknowledgements

We thank Jean-Christophe Ducom at the Scripps Research High Performance Computing for computational support, and Bill Anderson at the Scripps Research electron microscopy facility for microscope support. During this project D.G. was supported by a National Sciences Foundation predoctoral fellowship and G.C.L. was supported by the Pew Charitable Trusts as a Pew Scholar and an Amgen Young Investigator award. This project was also supported by the National Institutes of Health (NIH) grant DP2EB020402 to G.C.L. Computational analyses of EM data were performed on shared computing cluster funded by NIH grant S10OD021634 to G.C.L.

## Data Availability

Tilt series of microtubule-bound dynein-dynactin complexes are being uploaded to the Electron Microscopy Public Image Archive (EMPIAR) and will be made available as soon as possible.

